# Dissecting the role of glutamine in seeding peptide aggregation

**DOI:** 10.1101/2020.11.13.381632

**Authors:** Exequiel E. Barrera, Francesco Zonta, Sergio Pantano

## Abstract

Poly glutamine and glutamine-rich peptides play a central role in a plethora of pathological aggregation events. However, biophysical characterization of soluble oligomers —the most toxic species involved in these processes— remains elusive due to their structural heterogeneity and dynamical nature. Here, we exploit the high spatio-temporal resolution of simulations as a computational microscope to characterize the aggregation propensity and morphology of a series of polyglutamine and glutamine-rich peptides. Comparative analysis of ab-initio aggregation pinpointed a double role for glutamines. In the first phase, glutamines mediate seeding by pairing monomeric peptides, which serve as primers for higher-order nucleation. According to the glutamine content, these low molecular-weight oligomers may then proceed to create larger aggregates. Once within the aggregates, buried glutamines continue to play a role in their maturation by optimizing solvent-protected hydrogen bonds networks.

**TOC / Abstract Figure:** 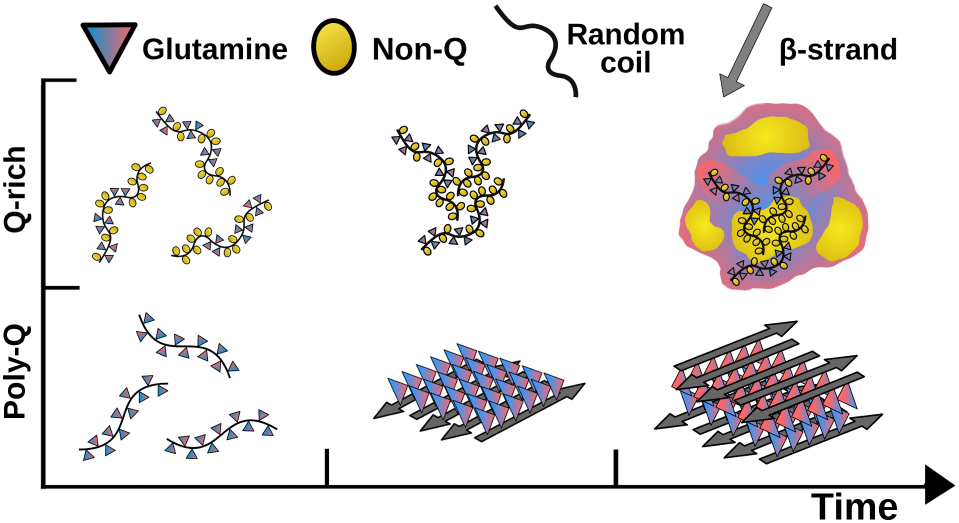

## INTRODUCTION

Protein misfolding, aggregation and their eponymous diseases have been vastly studied since the pioneering work of Alois Alzheimer in the beginnings of the twentieth century.^1^ After Alzheimer’s disease, multiple neurodegenerative diseases / protein dyads have been described, including frontotemporal dementia / TDP-43; Creutzfeldt-Jakob’s / prion-protein or Parkinson’s / α-synuclein. A special neurodegenerative subcategory is represented by the inherited polyglutamine diseases, where the neurotoxicity correlates with the length of consecutive glutamine tracts in the corresponding proteins. Here the dyads encompass Huntington’s / huntingtin; Kennedy’s / androgen receptor; spinocerebellar ataxia / ataxin-1, and dentatorubral-pallidoluysian atrophy / atrophin-1, among others. Abnormal protein aggregation is also present in systemic diseases like in the amyloid cardiomyopathy / transthyretin and type-2 diabetes / islet amyloid polypeptide. In the vastly studied Alzheimer’s amyloid cascade hypothesis, published by Hardy et al. in 1992, the key event proposed to trigger neurotoxicity is the formation of insoluble protein aggregates.^2^ However, Lambert et al. changed this paradigm, in 1998, shifting the investigation focus towards soluble oligomers as the main responsible for inhibition of long-term synaptic plasticity.^3^ Since then, all the above-mentioned protein aggregation diseases have been associated with these soluble, small aggregated species.^4–12^

The experimental structural determination of protein oligomeric species is extremely challenging because of their transient nature and structural heterogeneity.^13^ To cope with these difficulties, many computational studies have addressed the interaction modes of numerous aggregating peptides, including polyglutamine peptides, providing useful mechanistic insight.^14–21^ However, a comprehensive picture of the role of glutamine residues in different contexts is still missing. Among other reasons, this void originates in the substantial computational cost of atomically detailed simulations. This problem rapidly upscales, as different peptide sequences, multiple copies, and conditions are necessary to obtain a proper generalization in biologically relevant timescales.

We circumvent these limitations by employing cutting-edge coarse-grained simulations^22^ to examine glutamine’s role in early aggregation events. We studied a series of polyglutamine (poly-Q) and glutamine-rich (Q-rich) peptides displaying self-aggregative behavior.

A comparative analysis among homogeneous poly-Qs of different lengths and heterogeneous Q-rich peptides shows that glutamines play a double role in seeding aggregation. First, they participate in initial inter-monomer contacts, to then become the predominant residue mediating oligomer association by forming solvent-buried Hydrogen bonds.

Taken as a whole, the homogeneous set of simulations presented here offer original insight into the mechanistic role of glutamine in the initiation of pathogenic aggregation. Complementary, we present highly detailed scale and time resolution results that validate mechanistic pathways previously proposed by low-resolution experimental techniques.^23,24^

## RESULTS

Aimed to establish a comparative baseline for the aggregation of Q-rich peptides, we first investigate homogeneous poly-Q, progressively increasing the length of the peptides. To this aim we consider three different peptide lengths: Q4, Q11, and Q36.

### Tetraglutamine peptide (Q4)

Q4 in solution showed a quick transition from their initial, fully extended conformation towards a broad backbone-angle distribution, scattered on the fourth quadrant of the Ramachandran plot (**Supplementary figure 1A**). Along the time, Q4 peptides showed a continuous interchange between monomeric, and short-lived oligomeric species with a mean cluster size (MCS) of 1.94 ± 0.1 (**Figure 1A**). MCS can adopt values ranging from 1 (when all peptides in solution are monomeric species) up to the total number of simulated peptides (27, for a full aggregation in the Q4 case; see Methods). As seen from **Figure 1B**, monomeric species are dominant in solution, followed by dimers and trimers. Analysis of the global secondary structure (**Figure 1C**) evidenced only a small amount of extended and isolated β-structures, as a result of the fleeting character of the small clusters observed during the simulation (see molecular drawings in **Figure 1C**).

**Figure 1.**
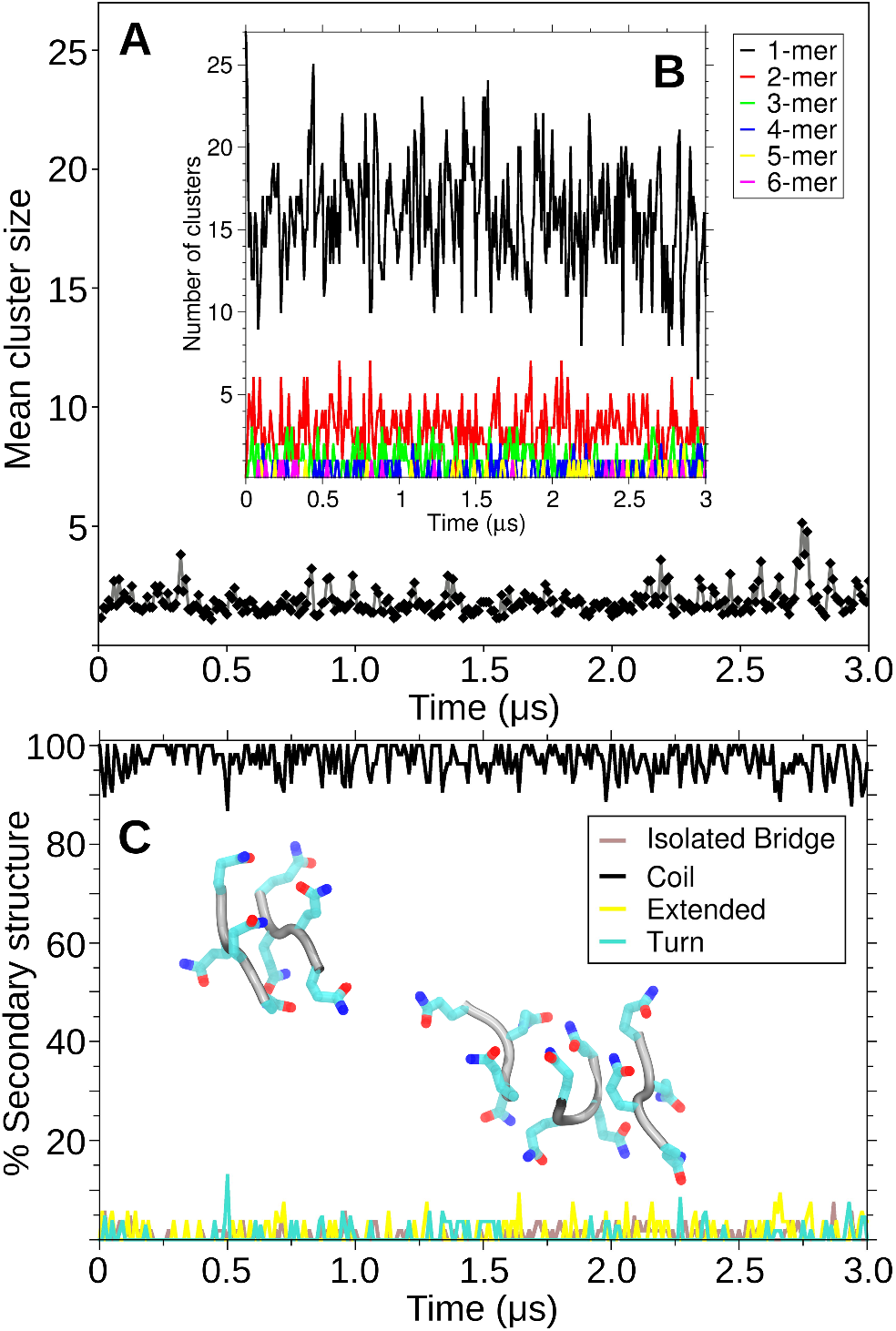
Aggregation analysis for Q4 peptides. (A) Mean cluster size measured throughout 3 μs MD simulations. (B) Number of clusters classified by size. (C) Global percentage secondary structure. Representative structures of Q4 dimers and trimers are drawn with a cartoon representation. Side chains are colored by element.

### Polyglutamine peptide Q11

Compared with the Q4 system, the longer Q11 peptide explored wider areas of the Ramachandran plot (**Supplementary figure 1B**). However, its length was insufficient to generate stable β hairpins-like motifs, and Q11 sampled mostly linear configurations. In contrast with the previous case, the MCS calculation indicates that the interaction between Q11 occurs in two well-differentiated fashions (**Figure 2A**). Initially, random encounters led to the formation of dimers or trimers (**Figure 2B**), which then served as nucleation seeds. Subsequently, all Q11 peptides cooperatively associate within a short time window. The dynamics of this process lead to the MCS’s saturation, aggregating all the peptides available in the simulation box. The progressive association of Q11 is accompanied by the passage from a random coil to parallel or antiparallel β-sheet conformations (**Figure 2C**). Indeed, the time series of the secondary structure for coil and β-sheets shows a scissors-like graph with a crossing point corresponding to the region where the associations become cooperative (**Figure 2D**). In the final state, almost 80% of the peptides are in β-sheet conformation, i.e., only 1-2 amino acids at the extremities of each peptide remain disordered.

**Figure 2.**
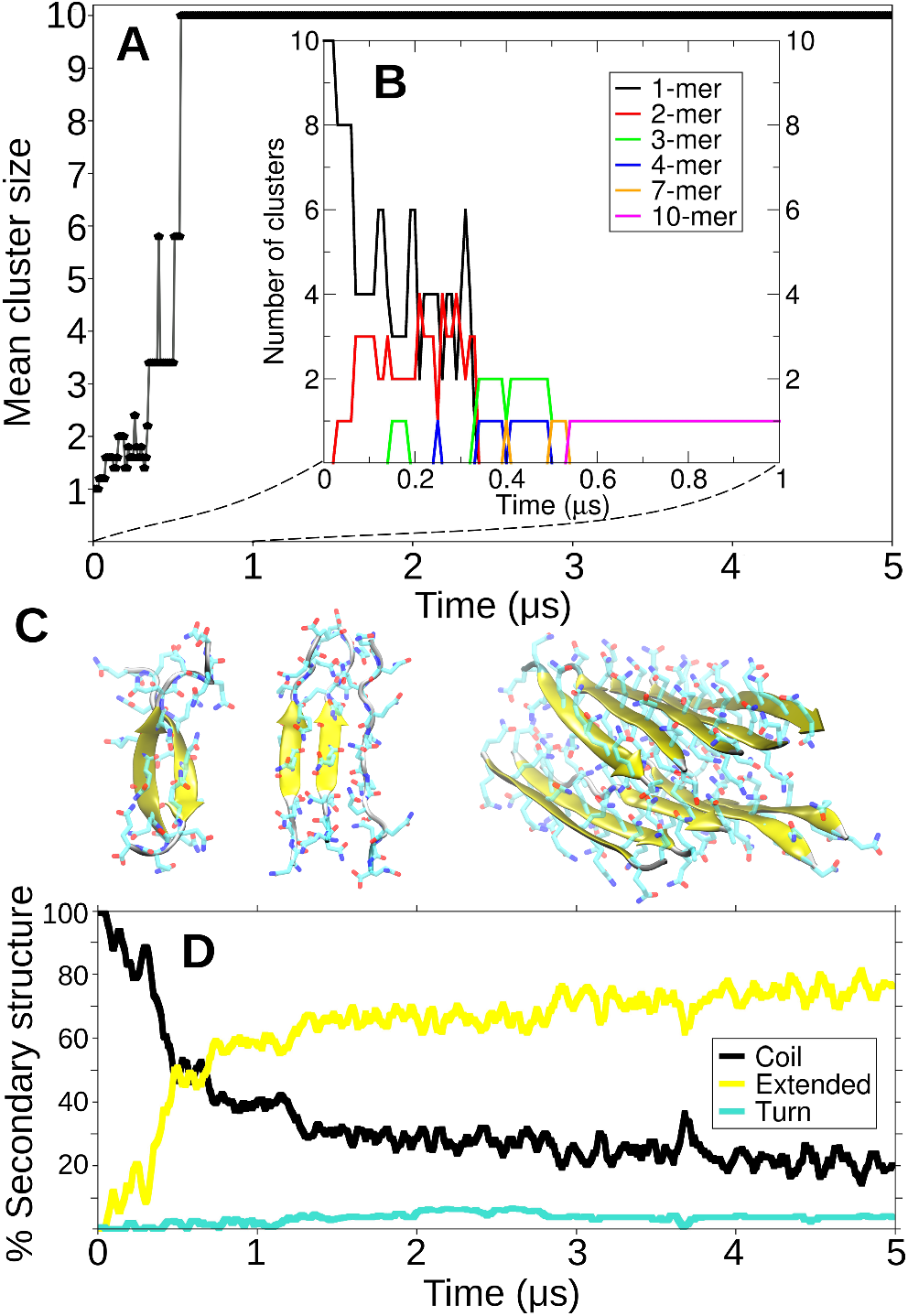
Aggregation of Q11 peptides. (A) Mean cluster size measured throughout 5 μs of MD simulations. (B) Number of clusters classified by size. The analysis is restricted to the first μs of the trajectory where all the association-dissociation events occur. (C) Cartoon representation of stable oligomers found for Q11. (D) Time evolution of the percentage of secondary structure content. The time scales in panels A and D are coincident.

### Polyglutamine peptide Q36

Increasing the polyglutamine size to 36 amino acids (Q36) conferred the poly-Q the capability to adopt turn conformations and forming β-hairpins. The accessibility to a wider conformational variety translates into different interaction possibilities. Despite this, in all aggregation events monitored, monomers always interacted with a pre-formed ß-stranded region in a neighboring molecule (**Figure 3A**). In close analogy with the Q11 case, after these association events, the β-sheet content increased reaching average values of 50.37% ± 5.32. The Q36 aggregates present intricate structural motifs, showing turn percentages of 9.81% ± 2.5 (**Figure 3** and **Supplementary figure 3A**).

**Figure 3.**
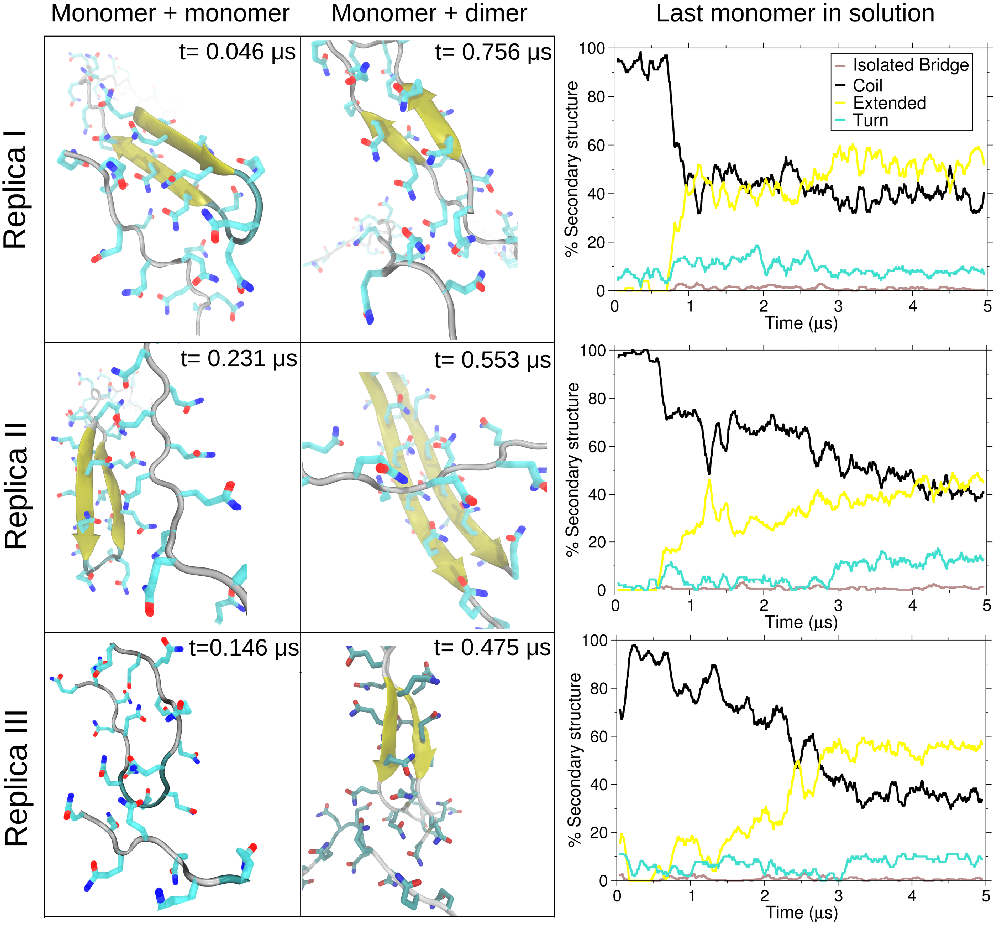
Aggregation of Q36 peptides. Snapshots of the association events for each simulated replica are shown in the left grid. Q36 monomers first contacts always involved interactions with preformed double stranded regions of their counterparts. Secondary structure plots are shown in the right panels.

### Q-Rich peptides

The results presented in the previous section point out that despite the well-known aggregation propensity of poly-Q peptides, there is a threshold below which, they can not sustain large scale aggregation. Therefore, we sought to characterize how both the glutamine content and the flanking regions of Q-rich peptides can affect their aggregative behavior.

### Alpha-gliadin peptide (p31-43)

As the first example of Q-rich peptide, we choose a derivative of the human protein alpha-gliadin. The peptide named p31-43 carries five glutamines in 13 amino acids (middle panel, **Figure 4A**) and is a proteolytic, gluten derived, peptide generated in the stomach and related to celiac disease.^25^ Using biophysical techniques, we showed that this peptide undergoes spontaneous aggregation with a concomitant but limited increase in the β-strand content.^26^

**Figure 4.**
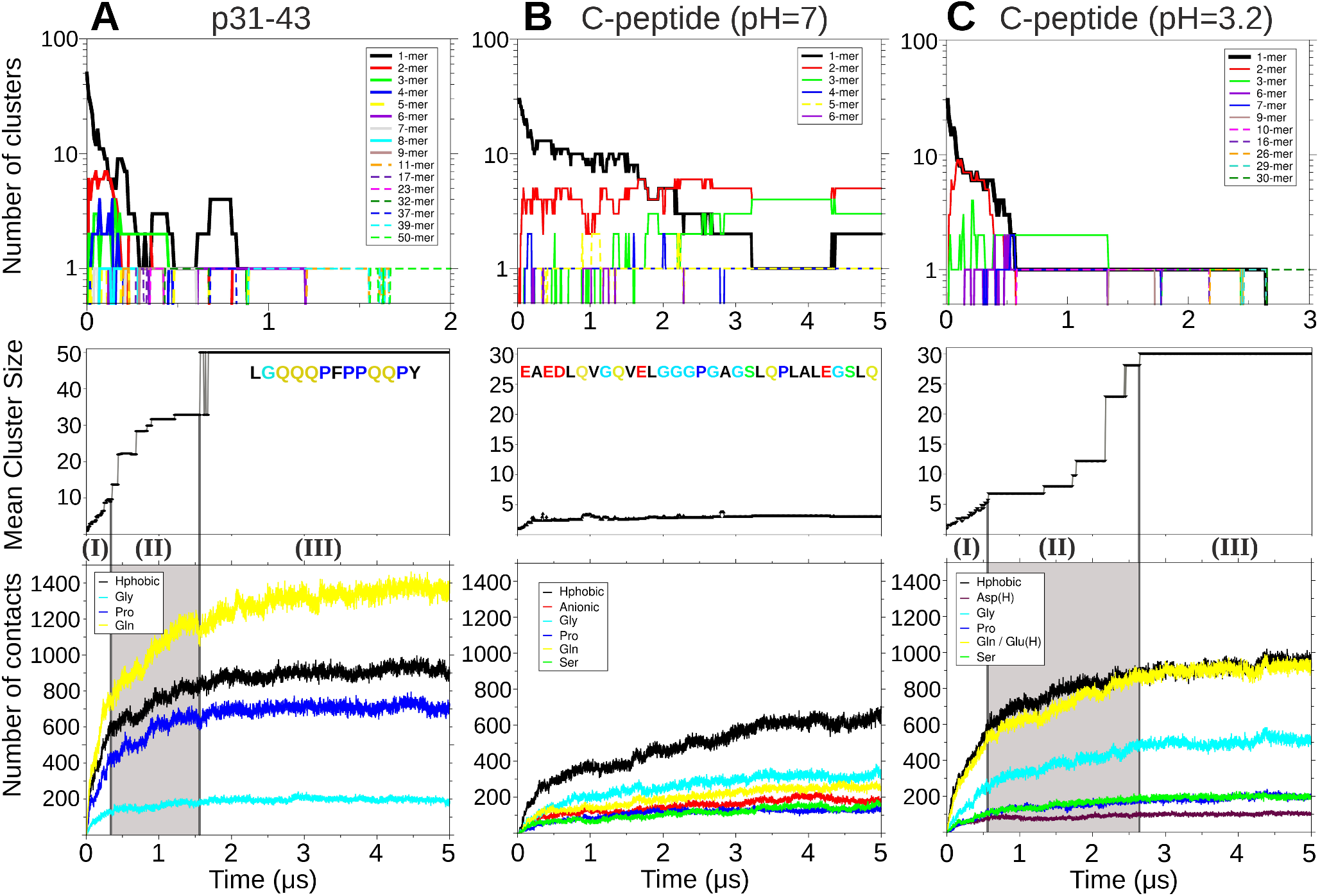
Aggregation analysis for the p31-43 peptide (A), C-peptide at pH=7 (B) and C-peptide at pH=3.2 (C). Top panels: Number of clusters classified by size, notice that the y axis is represented in a logarithmic scale. For (A) and (B) The analysis is focused on the first μs of simulation where most of the association events occur. Middle panels: Mean cluster size is measured throughout 5 μs of MD simulations. For (A) and (B) vertical lines divide the graph in three phases, according to the aggregation rate. The aminoacidic sequence of p31-43 and C-peptide is here shown. Lower panels: Number of interchain contacts defined by residue. Leucine, phenylalanine and tyrosine contacts are grouped as hydrophobic; aspartate and glutamate as anionic.

Here, to focus on glutamine, we simulated an ensemble of non-zwitterionic p31-43 and monitored the contribution of different amino acids during the aggregation process.

In contrast with poly-Q peptides, heterogeneous peptide aggregation follows different dynamics. According to the aggregation rate, we divide the analysis in three different phases. On phase I, monomers rapidly associate to form low molecular-weight oligomers (top panel, **Figure 4A**). This is followed by a slower aggregation (phase II), where oligomers progressively interact with each other until reaching a final amorphous 50-mer aggregate, presenting a radius of gyration of 2.48 ± 0.01 nm. The global content of β-extended structures raises from zero to 28.08 % ± 1.71. The cluster size remains stable until the end of the trajectory (phase III). A per-residue interpeptide contact analysis (lower panel, **Figure 4A**) indicates that glutamines lead the p31-43 aggregation process, followed by hydrophobic residues (Leu, Phe, and Tyr), proline, and glycine, to a much lesser extent. This is in line with the relative amino acid abundances in this peptide). If we divide the contact analysis in the three above mentioned aggregation phases we can observe how glutamine-mediated contacts are increased in phase II (gray shaded time window in lower panel of **Figure 4A** and **Supplementary video 1**). In phase III, contacts established by all amino acids reach a plateau, except for glutamine. Therefore, not only glutamines mediate oligomer association but they also continue optimizing the formation of Hydrogen bonds after all peptides are clustered, suggesting a sort of maturation within the final aggregated state.

### Proinsulin’s connecting peptide (C-peptide)

Aimed to further explore glutamines’ role in an entirely unrelated system, we focus on insulin’s C-peptide, which undergoes only limited aggregation. At pH=7, the C-peptide presents monomers and various low molecular weight oligomers compatible with dimers to pentamers.^27^ C-peptide is 31 amino acids long with only four —two of them are highly conserved— glutamines^28^ (middle panel, **Figure 4B**). However, it also contains four conserved glutamic acids, and at pH=3.2 has been shown to form oligomers and large amyloid-like aggregates with a high β-sheet content.^27^ Since at pH=3.2, all carboxyl groups are expected to be protonated,^29^ its Hydrogen-bonding donor-acceptor capabilities could resemble those present in glutamine. Although this might seem a chemically naïve approximation, quantum calculations yielded similar Hydrogen bond energies to protonated carboxyl and amide groups.^30^ The same substitution has been proven as a conservative mutation in a glutamate transporter with a deeply buried acidic residue.^31^ Hence, at low pH values, we will consider C-peptide as a Q-rich molecule.

Not surprisingly, the aggregation dynamics and kinetics of the C-peptide at pH=7 differ from those seen in the previous systems (**Figure 4B**). Unlike in the other cases, the MCS converges to nearly 3, even though there is a ten fold excess of free peptides in the solution. On a first phase the monomer population rapidly decreases, forming multiple low molecular weight oligomers (top panel, **Figure 4B**). After this period the aggregation rate diminishes, presenting a monomer-dimer-trimer equilibrium^32^, and the presence of two stable tetrameric and pentameric species. A per-residue analysis clearly shows that the limited aggregation is driven by hydrophobic forces and most likely limited by the electrostatic repulsion conferred by the anionic residues (lower panel, **Figure 4B** and **Supplementary video 2**).

Converting the C-peptide into Q-rich like peptide by decreasing the solution’s pH recovered an aggregation-prone behavior. This time a globular 30-mer aggregate was formed, with a radius of gyration of 3.78 ± 0.06 nm and 15.85 % ± 0.84 of β-extended structures. Although with slower kinetics, the evolution of contacts per residue displayed a behavior similar to p31-43, with an initial depletion of monomers forming dimers, trimers and hexamers (top panel, **Figure 4C**). In close analogy with p31-43, oligomers undergo progressive association during phase II, until one single aggregate is formed. As can be seen in the lower panel of **Figure 4C,** phase I is mainly driven by hydrophobic contacts and hydrogen bonding between glutamines or protonated glutamic acids in proportions according to their abundances. During aggregation phase II, this contact profile is overturned with Hydrogen bonds contributing the most to oligomer association as can be seen in the steepest evolution of Q-like contacts. These glutamine mediated contacts also participate in the compaction of the final aggregate, as can be seen in **Supplementary video 3**.

## DISCUSSION AND CONCLUSIONS

Our analysis of the role of glutamine within different peptide contexts revealed some general features summarized in **Figure 5**.

**Figure 5.**
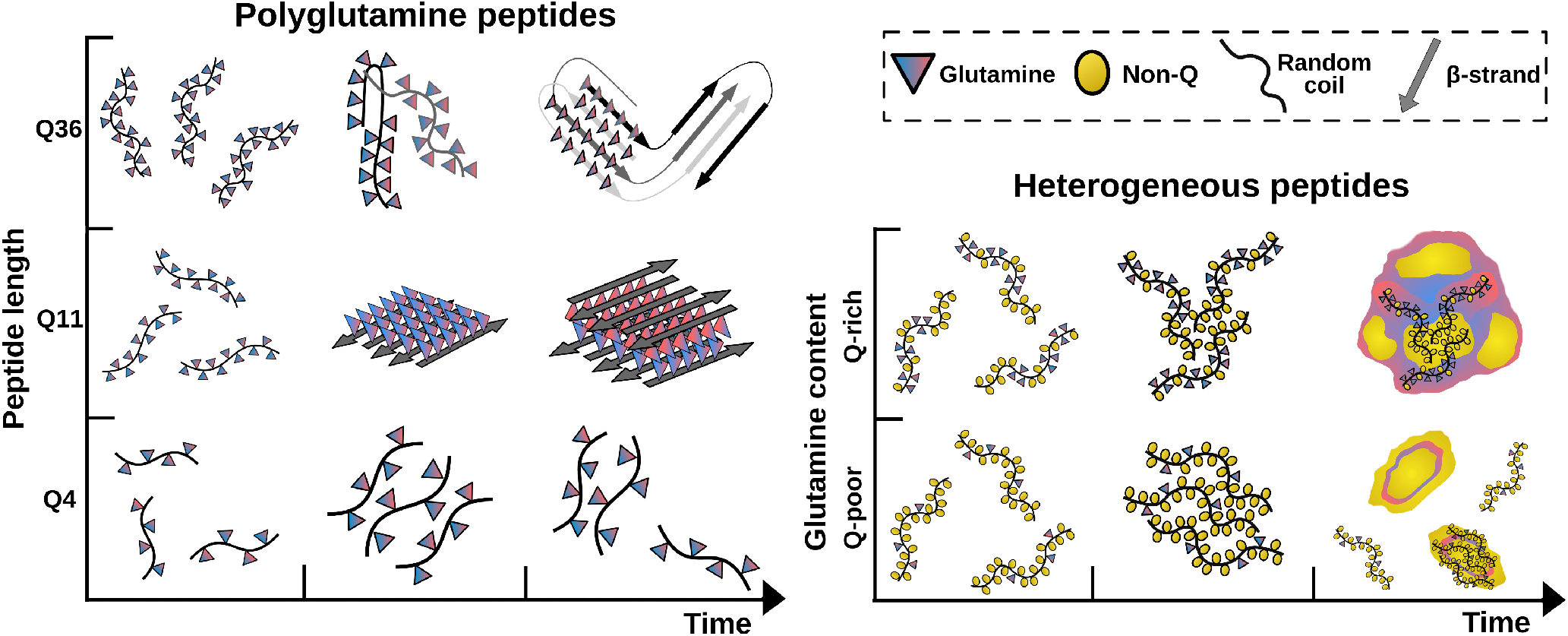
Schematic description of the different aggregation scenarios studied. The time evolution of the aggregation is classified by peptide length for poly-Q peptides (left grid) and by glutamine content for heterogeneous peptides (right grid).

In the case of poly-Q, the peptide length affects both the extent and the morphology of aggregates. Q4 peptides show high solubility, presenting rapid inter-conversions between monomeric, dimeric, and trimeric species (**Figure 1B**). However, it is widely accepted that expanding glutamine tracts increase their aggregation propensity, particularly enriching their β-sheet contents.^33^ Therefore, we decided to study two longer poly-Q peptides: Q11 and Q36, representing expansions below and above the toxic threshold in Huntington’s disease.^34^ Q11 peptides initially form dimers, trimers, and tetramers stabilized by multiple backbone hydrogen bonds, that ultimately result in the recruitment of all peptides in the solution. In line with previous studies, the β-sheets produced present extensive surfaces with side chains alternating Hydrogen bonds donors and acceptors that recruit new peptides to end in a putative crystalline β-sheet structure.^35^ In contrast, Q36 presents higher conformational flexibility, allowing β-hairpin formation. These double-stranded motifs acted as aggregation-prone regions. These regions interact with unstructured soluble monomers and promote a conformational transition towards β-sheet rich structures, resembling a nucleated-elongation model.^23^ This mechanism has been previously proposed by Wetzel et al. studying different length poly-Q peptides using a microtiter plate elongation assay.^36^ The distinctive structural characteristics that we observed in poly-Q peptides of different lengths are also in remarkable agreement with experimental data obtained by IR spectroscopy, validating the length-dependent aggregation pathways proposed in this low-resolution structural determination.^24^

In contrast with homogeneous poly-Qs, for heterogeneous glutamine peptides, the relevant variable is their relative aminoacidic abundances. Analyzing its content in relative terms provides a convenient and quantitative characterization. To this aim, we calculated the ratio between glutamine and non-Q amino acids and compared them with the ratio of new intermolecular contact established by glutamines vs non-Q amino acids, in each aggregation phase.

The C-peptide at pH=7 displays a glutamine ratio of only 0.15 (**Table 1**). In this Q-poor peptide, glutamine contacts are not dominant and remain insensitive to the aggregation phase. It is somehow predictable that progressively increasing glutamine content could increase their engagement in interpeptide contacts. However, the remarkable observation is how these interactions become dominant during the association of low molecular weight oligomers (phase II), with contact ratios surpassing residue fraction (gray shadowed row, **Table 1**). Calculating relative contents, this time between glutamine and hydrophobic amino acids, indicate that initial monomeric association is ruled by the hydrophobic effect and is later switched to a Q-driven regime.

**Table 1.** Glutamine abundance and contact ratios for the Q-poor and Q-rich systems. Contacts ratios are calculated separately for each aggregation phase. The values are also reported relative to hydrophobic residues (Q/HP).

|  | C-peptide (pH = 7) |  | C-peptide (pH = 3.2) |  | p31-43 |  |
| --- | --- | --- | --- | --- | --- | --- |
|  | Q/Non-Q | Q/HP | Q/Non-Q | Q/HP | Q/Non-Q | Q/HP |
| Residue abundance | 0.15 | 0.36 | 0.35 | 0.73 | 0.63 | 1.67 |
| Contacts phase I | 0.20 | 0.42 | 0.45 | 0.89 | 0.66 | 1.27 |
| Contacts phase II | 0.15 | 0.34 | 0.53 | 1.21 | 0.82 | 1.79 |
| Contacts phase III |  | - | 0.43 | 0.78 | 1.21 | 2.33 |

In a nutshell, the extension of poly-Q peptides is determinant for their aggregation, showing three well-differentiated behaviors. Only above a certain length, poly-Qs may display highly ordered (nearly crystalline) aggregation, after which the possibility to form intramolecular hairpins may lead to large scale but conformationally heterogeneous fibrillar arrays (**Figure 5**-left side and **Supplementary Figure 3A**). In contrast to poly-Q, heterogeneous peptides present as many variables as the amino acidic composition can introduce. In this work, we focused on glutamine content and showed how its increase could trigger the low molecular weight oligomers’ association, stabilizing large globular aggregates (**Figure 5**-right side and **Supplementary Figure 3B-C**).

Taken as a whole, our data propose that an early blocking of the glutamine-mediated interactions could impede aggregation, shifting the equilibrium from low molecular weight oligomers to monomeric species. This could facilitate clearance mechanisms like those observed in huntingtin and its autophagic degradation that acts on soluble but not preformed aggregates.^37^

Several efforts have been devoted to design short peptides with the capability to modulate toxic protein aggregation.^38,39^ Just as an example, we explored the capacity of two pentapeptides to modulate Q11 aggregation. For this, we test non-zwitterionic Q5 and QEQQQ. Compared with the homogeneous Q11 system, both pentapetides alter the aggregation’s kinetics (**Supplementary Figure 4A**), although the content of β-extended conformations converged to indistinguishable values (**Supplementary Figure 4B**). However, the pentapeptides’ presence, especially QEQQQ, modified the aggregate’s final topologies (**Supplementary Figure 4C-E**). The procedure presented here provides a robust and costeffective computational strategy to test for different peptides’ aggregation propensity or explore conditions that may modify these propensities

## METHODS

We generate peptides’ initial coordinates with Chimera^40^ visualization tool, setting the backbone torsional angles as fully extended conformations. C-peptide was studied at pH values of 3.2 and 7, defining their protonation states with PropKa.^41^ Atomic structures were mapped to CG beads using the map files included in SIRAH Tools.^42^ Carboxy- and amino-terminals were set as neutral, except for the C-peptide where we focused on the effects of electrostatic interactions over aggregation properties; in these systems terminals were set as zwitterionic. MD simulations were performed by triplicate with GROMACS 2018.4.^43^ All the six peptides were centered in cubic boxes, which sizes were defined setting a distance of 1.5 nm between the solute and the box boundaries. Systems were solvated using a pre-equilibrated box of SIRAH’s water model (named WT4).^44^ Forces on the CG beads were balanced applying 5000 iterations of the steepest descent algorithm. The heating step was performed using the V-rescale thermostat,^45^ keeping the pressure at 1 bar with the Parrinello-Rahman barostat.^46^ In order to generate initial conformations for the aggregation studies, 1 μs production runs were simulated, choosing different conformers from the last 0.1 μs of trajectory. Multiple copies of these conformers were placed in cubic boxes setting a distance of 4 nm between their geometric centers. An identical system setup as above described for the monomeric peptides, was applied for the multiple-peptide simulation systems. MD trajectories’ analysis included secondary structure determinations. To achieve this we first employed the backmapping utility of SIRAH tools and then assign secondary contents to the reconstructed atomic coordinates with STRIDE.^47^ To estimate the size of the aggregates along the simulations we calculated the Mean Cluster Size (MCS) as it has been previously defined by Taiji et. Al^48^: 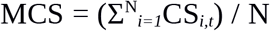. “N” corresponds to the total number of peptides in the simulation box and “CS” is the number of peptides forming a cluster which peptide “*i*” belongs at time “*t*”. The generation of a cluster was defined using a distance cutoff of 6 Å between beads of different peptides. Interpeptide contacts were measured with the GROMACS utility *gmx mindist* and the MCS calculation was performed with an *in house* Python script. The number of total interpeptide contacts defined by residue was calculated with the Tcl script: *newcontacts.tcl* downloaded from https://www.ks.uiuc.edu/. Here, to better discriminate the residues involved in interpeptide contacts, we used a smaller cutoff value of 5 Å.

## ACKNOWLEDGMENTS

This work was partially funded by FOCEM (MERCOSUR Structural Convergence Fund), COF 03/11 and the National Natural Science Foundation of China (NSFC grant 31770776 to FZ). EEB and SP belong to the SNI program of ANII. EEB is a beneficiary of a postdoctoral fellowship of CONICET. Simulations were performed on the National Uruguayan Center for Supercomputing, ClusterUY. We thank Engr. Martin Etchart for his valuable collaboration in Python scripting.

## SUPPLEMENTARY INFORMATION

**Supplementary figure 1.**
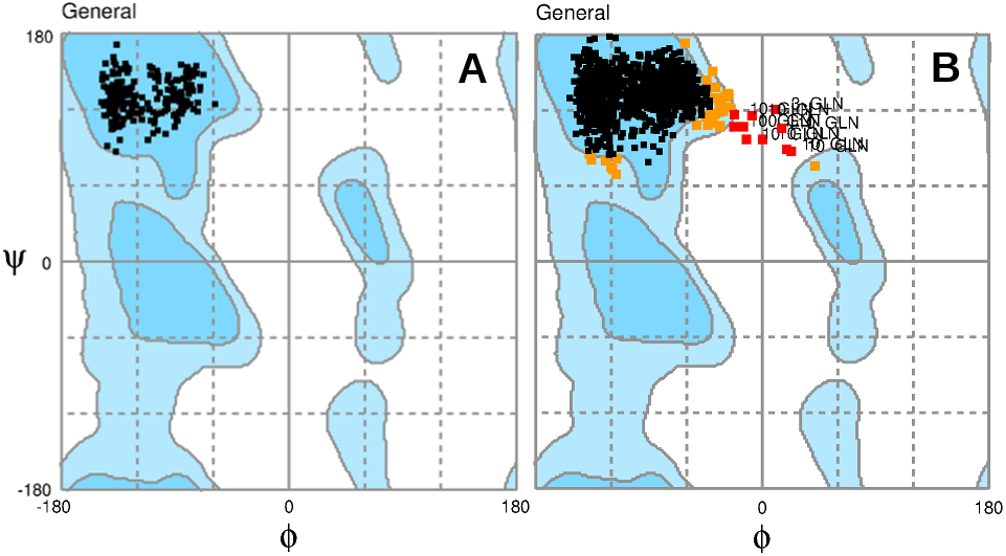
Ramachandran plots for Q4 (A) and Q11 peptides (B). Black, orange and red dots correspond to most favored, allowed, and less favored regions of the map, respectively.

**Supplementary figure 2.**
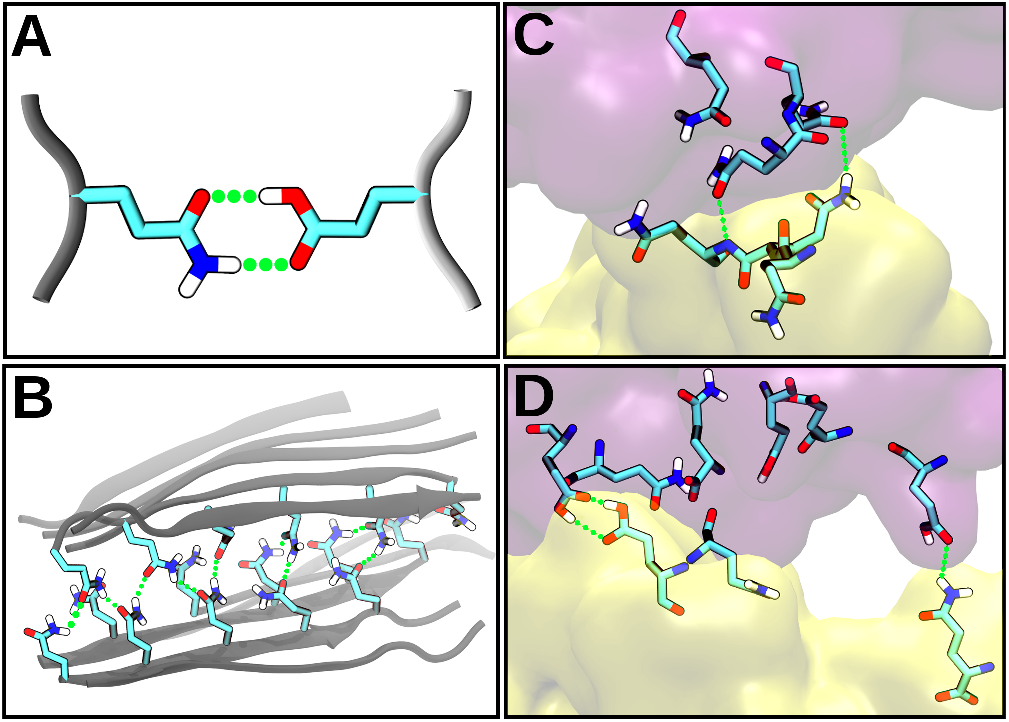
(A) Schematic representation of a double Hydrogen bond (green dotted lines) between glutamine and a protonated glutamic acid. (B) Two β-sheets of Q11 peptides stabilized by inter side-chain H-bonds. (C) Interface between two p31-43 oligomers mediated by glutamine H-bonds. (D) Same as C for glutamines and protonated glutamic acids in a C-peptide aggregate at pH=3.2.

**Supplementary Figure 3.**
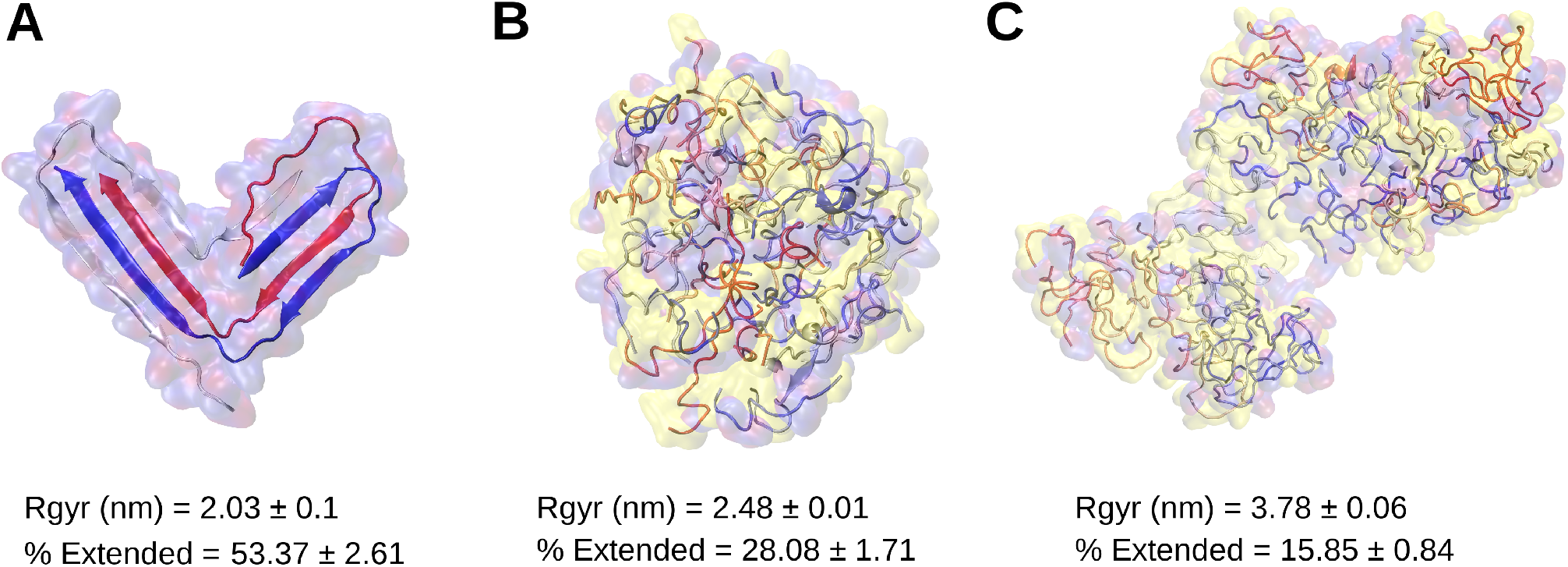
Final snapshots from 5 μs trajectories for (A) Q36, (B) p31-43 and (C) C-peptide (pH=3.2). Peptide’s backbones are drawn in cartoon representations and colored by fragment. Van der Waals surfaces are transparent and colored red/blue (glutamine residues) and yellow (non-glutamine residues). Radius of gyration and β-extended conformations content are averaged over the last 0.1 μs of the corresponding trajectories.

**Supplementary Figure 4.**
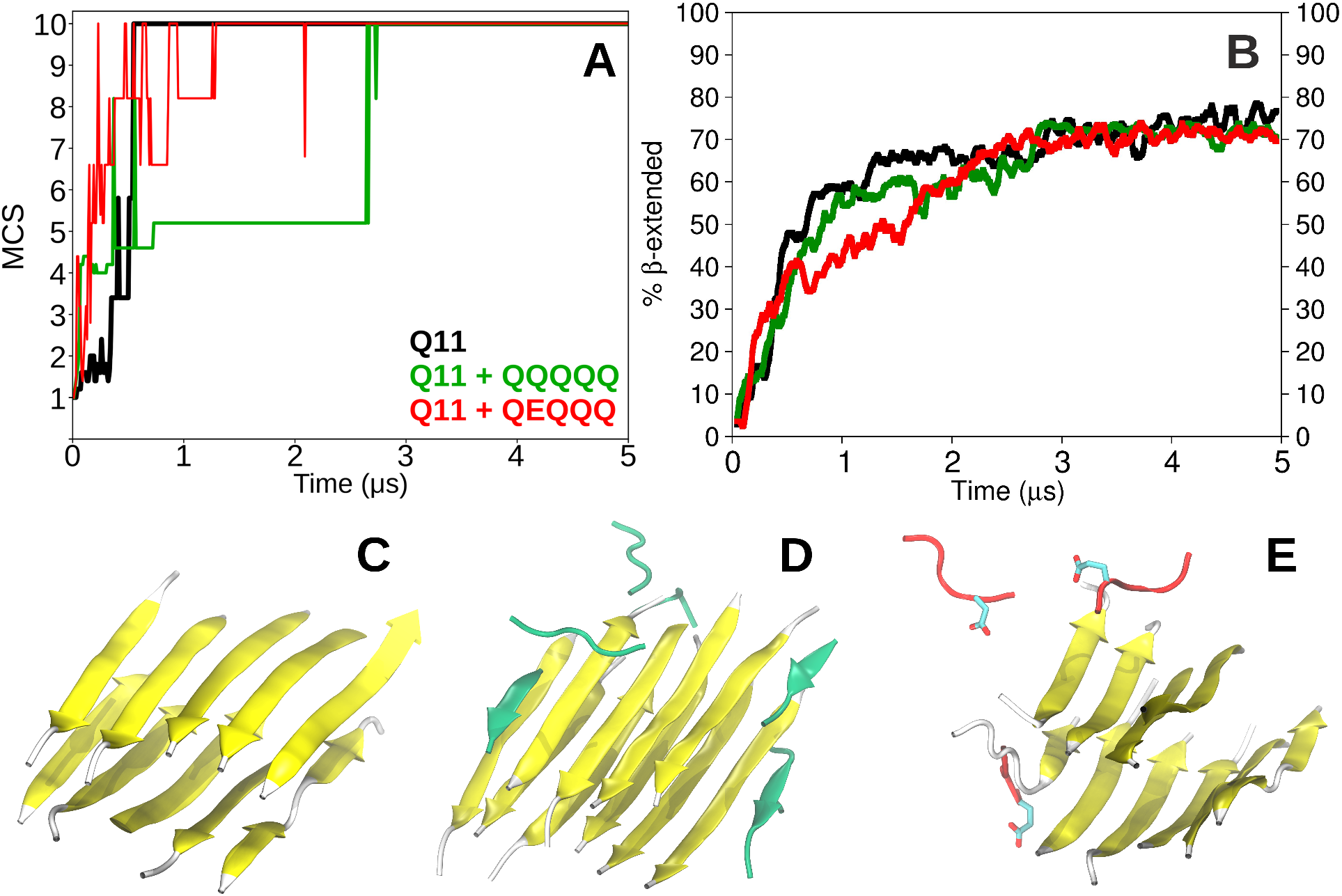
Modulation of Q11 aggregation exerted by different pentapeptides along 5 μs trajectories. Mean Cluster Size evolution (A) and β-extended percentage content (B) of Q11 in each of the three simulated systems. Cartoon representation of the final snapshots for the homogeneous Q11 (C), Q11 + Q5 (D) and Q11 + QEQQQ systems (E).

## REFERENCES

(1) Alzheimer; A. Uber Einen Eigenartigen Schweren Erkrankungsprozess Der Hirninde. Neurol. Cent. 1906, 25, 1134.

(2) Hardy, J. A.; Higgins, G. A. Alzheimer’s Disease: The Amyloid Cascade Hypothesis. Science. American Association for the Advancement of Science April 10, 1992, pp 184–185. https://doi.org/10.1126/science.1566067.

(3) Lambert, M. P.; Barlow, A. K.; Chromy, B. A.; Edwards, C.; Freed, R.; Liosatos, M.; Morgan, T. E.; Rozovsky, I.; Trommer, B.; Viola, K. L.; et al. Diffusible, Nonfibrillar Ligands Derived from Aβ1-42 Are Potent Central Nervous System Neurotoxins. Proc. Natl. Acad. Sci. U. S. A. 1998, 95 (11), 6448–6453. https://doi.org/10.1073/pnas.95.11.6448.

(4) Fang, Y. S.; Tsai, K. J.; Chang, Y. J.; Kao, P.; Woods, R.; Kuo, P. H.; Wu, C. C.; Liao, J. Y.; Chou, S. C.; Lin, V.; et al. Full-Length TDP-43 Forms Toxic Amyloid Oligomers That Are Present in Frontotemporal Lobar Dementia-TDP Patients. Nat. Commun. 2014, 5 (1), 1–13. https://doi.org/10.1038/ncomms5824.

(5) Bengoa-Vergniory, N.; Roberts, R. F.; Wade-Martins, R.; Alegre-Abarrategui, J. Alpha-Synuclein Oligomers: A New Hope. Acta Neuropathologica. Springer Verlag December 1, 2017, pp 819–838. https://doi.org/10.1007/s00401-017-1755-1.

(6) Silveira, J. R.; Raymond, G. J.; Hughson, A. G.; Race, R. E.; Sim, V. L.; Hayes, S. F.; Caughey, B. The Most Infectious Prion Protein Particles. Nature 2005, 437 (7056), 257–261. https://doi.org/10.1038/nature03989.

(7) Beitel, L. K.; Alvarado, C.; Mokhtar, S.; Paliouras, M.; Trifiro, M. Mechanisms Mediating Spinal and Bulbar Muscular Atrophy: Investigations into Polyglutamine-Expanded Androgen Receptor Function and Dysfunction. Front. Neurol. 2013, 4 (53). https://doi.org/10.3389/fneur.2013.00053.

(8) Lasagna-Reeves, C. A.; Rousseaux, M. W. C.; Guerrero-Munoz, M. J.; Vilanova-Velez, L.; Park, J.; See, L.; Jafar-Nejad, P.; Richman, R.; Orr, H. T.; Kayed, R.; et al. Ataxin-1 Oligomers Induce Local Spread of Pathology and Decreasing Them by Passive Immunization Slows Spinocerebellar Ataxia Type 1 Phenotypes. Elife 2015, 4 (DECEMBER 2015), 1–11. https://doi.org/10.7554/eLife.10891.

(9) Schonhoft, J. D.; Monteiro, C.; Plate, L.; Eisele, Y. S.; Kelly, J. M.; Boland, D.; Parker, C. G.; Cravatt, B. F.; Teruya, S.; Helmke, S.; et al. Peptide Probes Detect Misfolded Transthyretin Oligomers in Plasma of Hereditary Amyloidosis Patients. Sci. Transl. Med. 2017, 9 (407). https://doi.org/10.1126/scitranslmed.aam7621.

(10) Haataja, L.; Gurlo, T.; Huang, C. J.; Butler, P. C. Islet Amyloid in Type 2 Diabetes, and the Toxic Oligomer Hypothesis. Endocrine Reviews. The Endocrine Society May 2008, pp 303–316. https://doi.org/10.1210/er.2007-0037.

(11) Kim, Y. E.; Hosp, F.; Frottin, F.; Ge, H.; Mann, M.; Hayer-Hartl, M.; Hartl, F. U. Soluble Oligomers of PolyQ-Expanded Huntingtin Target a Multiplicity of Key Cellular Factors. Mol. Cell 2016, 63 (6), 951–964. https://doi.org/10.1016/j.molcel.2016.07.022.

(12) Leitman, J.; Ulrich Hartl, F.; Lederkremer, G. Z. Soluble Forms of PolyQ-Expanded Huntingtin Rather than Large Aggregates Cause Endoplasmic Reticulum Stress. Nat. Commun. 2013, 4, 1–10. https://doi.org/10.1038/ncomms3753.

(13) Tycko, R. Molecular Structure of Aggregated Amyloid-β: Insights from Solid State Nuclear Magnetic Resonance. Cold Spring Harb. Perspect. Med. 2016, 6 (8), a024083. https://doi.org/10.1101/cshperspect.a024083.

(14) Nguyen, P. H.; Sterpone, F.; Derreumaux, P. Aggregation of Disease-Related Peptides, 1st ed.; Elsevier Inc., 2020. https://doi.org/10.1016/bs.pmbts.2019.12.002.

(15) Morriss-Andrews, A.; Shea, J. E. Computational Studies of Protein Aggregation: Methods and Applications. Annu. Rev. Phys. Chem. 2015, 66 (January), 643–666. https://doi.org/10.1146/annurev-physchem-040513-103738.

(16) Carballo-Pacheco, M.; Ismail, A. E.; Strodel, B. On the Applicability of Force Fields to Study the Aggregation of Amyloidogenic Peptides Using Molecular Dynamics Simulations. J. Chem. Theory Comput. 2018, 14 (11), 6063–6075. https://doi.org/10.1021/acs.jctc.8b00579.

(17) Ruff, K. M.; Khan, S. J.; Pappu, R. V. A Coarse-Grained Model for Polyglutamine Aggregation Modulated by Amphipathic Flanking Sequences. Biophys. J. 2014, 107 (5), 1226–1235. https://doi.org/10.1016/j.bpj.2014.07.019.

(18) Haaga, J.; Gunton, J. D.; Buckles, C. N.; Rickman, J. M. Early Stage Aggregation of a Coarse-Grained Model of Polyglutamine. J. Chem. Phys. 2018, 148 (4). https://doi.org/10.1063/1.5010888.

(19) Fluitt, A. M.; De Pablo, J. J. An Analysis of Biomolecular Force Fields for Simulations of Polyglutamine in Solution. Biophys. J. 2015, 109 (5), 1009–1018. https://doi.org/10.1016/j.bpj.2015.07.018.

(20) Wang, Y.; Voth, G. A. Molecular Dynamics Simulations of Polyglutamine Aggregation Using Solvent-Free Multiscale Coarse-Grained Models. J. Phys. Chem. B 2010, 114 (26), 8735–8743. https://doi.org/10.1021/jp1007768.

(21) Flöck, D.; Rossetti, G.; Daidone, I.; Amadei, A.; Di Nola, A. Aggregation of Small Peptides Studied by Molecular Dynamics Simulations. Proteins Struct. Funct. Genet. 2006, 65 (4), 914–921. https://doi.org/10.1002/prot.21168.

(22) Machado, M. R.; Barrera, E. E.; Klein, F.; Sóñora, M.; Silva, S.; Pantano, S. The SIRAH 2.0 Force Field: Altius, Fortius, Citius. J. Chem. Theory Comput. 2019, 15 (4), 2719–2733. https://doi.org/10.1021/acs.jctc.9b00006.

(23) Bhattacharyya, A. M.; Thakur, A. K.; Wetzel, R. Polyglutamine Aggregation Nucleation: Thermodynamics of a Highly Unfavorable Protein Folding Reaction. Proc. Natl. Acad. Sci. U. S. A. 2005, 102 (43), 15400–15405. https://doi.org/10.1073/pnas.0501651102.

(24) Yushchenko, T.; Deuerling, E.; Hauser, K. Insights into the Aggregation Mechanism of PolyQ Proteins with Different Glutamine Repeat Lengths. Biophys. J. 2018, 114 (8), 1847–1857. https://doi.org/10.1016/j.bpj.2018.02.037.

(25) Gómez Castro, M. F.; Miculán, E.; Herrera, M. G.; Ruera, C.; Perez, F.; Prieto, E. D.; Barrera, E.; Pantano, S.; Carasi, P.; Chirdo, F. G. P31-43 Gliadin Peptide Forms Oligomers and Induces NLRP3 Inflammasome/Caspase 1-Dependent Mucosal Damage in Small Intestine. Front. Immunol. 2019, 10, 31. https://doi.org/10.3389/fimmu.2019.00031.

(26) Herrera, M. G.; Gómez Castro, M. F.; Prieto, E.; Barrera, E.; Dodero, V. I.; Pantano, S.; Chirdo, F. Structural Conformation and Self-Assembly Process of P31-43 Gliadin Peptide in Aqueous Solution. Implications for Celiac Disease. FEBS J. 2020, 287 (10), 2134–2149. https://doi.org/10.1111/febs.15109.

(27) Lind, J.; Lindahl, E.; Perálvarez-Marín, A.; Holmlund, A.; Jörnvall, H.; Mäler, L. Structural Features of Proinsulin C-Peptide Oligomeric and Amyloid States. FEBS J. 2010, 277 (18), 3759–3768. https://doi.org/10.1111/j.1742-4658.2010.07777.x.

(28) Munte, C. E.; Vilela, L.; Kalbitzer, H. R.; Garratt, R. C. Solution Structure of Human Proinsulin C-Peptide. FEBS J. 2005, 272 (16), 4284–4293. https://doi.org/10.1111/j.1742-4658.2005.04843.x.

(29) Kozlowski, L. P. IPC – Isoelectric Point Calculator. Biol. Direct 2016, 11 (1), 55–55. https://doi.org/10.1186/s13062-016-0159-9.

(30) Nie, B.; Stutzman, J.; Xie, A. A Vibrational Spectral Maker for Probing the Hydrogen-Bonding Status of Protonated Asp and Glu Residues. Biophys. J. 2005, 88 (4), 2833–2847. https://doi.org/10.1529/biophysj.104.047639.

(31) Mwaura, J.; Tao, Z.; James, H.; Albers, T.; Schwartz, A.; Grewer, C. Protonation State of a Conserved Acidic Amino Acid Involved in Na + Binding to the Glutamate Transporter EAAC1. ACS Chem. Neurosci. 2012, 3 (12), 1073–1083. https://doi.org/10.1021/cn300163p.

(32) Betz, S.; Fairman, R.; O’Neil, K.; Lear, J.; Degrado, W. Design of Two-Stranded and Three-Stranded Coiled-Coil Peptides. Philos. Trans. R. Soc. Lond. B. Biol. Sci. 1995, 348 (1323), 81–88. https://doi.org/10.1098/rstb.1995.0048.

(33) Petrakis, S.; Schaefer, M. H.; Wanker, E. E.; Andrade-Navarro, M. A. Aggregation of PolyQ-Extended Proteins Is Promoted by Interaction with Their Natural Coiled-Coil Partners. BioEssays 2013, 35 (6), 503–507. https://doi.org/10.1002/bies.201300001.

(34) Bates, G. P.; Dorsey, R.; Gusella, J. F.; Hayden, M. R.; Kay, C.; Leavitt, B. R.; Nance, M.; Ross, C. A.; Scahill, R. I.; Wetzel, R.; et al. Huntington Disease. Nat. Rev. Dis. Prim. 2015, 1 (April), 1–21. https://doi.org/10.1038/nrdp.2015.5.

(35) Viney, C. Natural Protein Fibers. Ref. Modul. Mater. Sci. Mater. Eng. 2016, 1–9. https://doi.org/10.1016/b978-0-12-803581-8.02271-2.

(36) Berthelier, V.; Wetzel, R. Screening for Modulators of Aggregation with a Microplate Elongation Assay. Methods Enzymol. 2006, 413 (06), 313–325. https://doi.org/10.1016/S0076-6879(06)13016-5.

(37) Hegde, R. N.; Chiki, A.; Petricca, L.; Martufi, P.; Arbez, N.; Mouchiroud, L.; Auwerx, J.; Landles, C.; Bates, G. P.; Singh-Bains, M. K.; et al. TBK1 Phosphorylates Mutant Huntingtin and Suppresses Its Aggregation and Toxicity in Huntington’s Disease Models. EMBO J. 2020, 39 (17). https://doi.org/10.15252/embj.2020104671.

(38) He, R. Y.; Lai, X. M.; Sun, C. S.; Kung, T. S.; Hong, J. Y.; Jheng, Y. S.; Liao, W. N.; Chen, J. K.; Liao, Y. F.; Tu, P. H.; et al. Nanoscopic Insights of Amphiphilic Peptide against the Oligomer Assembly Process to Treat Huntington’s Disease. Adv. Sci. 2020, 7 (2). https://doi.org/10.1002/advs.201901165.

(39) Diao, J.; Yang, K.; Lai, Y.; Li, Y.; Jiang, L.; Li, D.; Lu, J.; Cao, Q.; Wang, C.; Zheng, J.; et al. Structure-Based Peptide Inhibitor Design of Amyloid-β Aggregation. 2019. https://doi.org/10.3389/fnmol.2019.00054.

(40) Pettersen, E. F.; Goddard, T. D.; Huang, C. C.; Couch, G. S.; Greenblatt, D. M.; Meng, E. C.; Ferrin, T. E. UCSF Chimera. A Visualization System for Exploratory Research and Analysis. J. Comput. Chem. 2004, 25 (13), 1605–1612. https://doi.org/10.1002/jcc.20084.

(41) Rostkowski, M.; Olsson, M. H.; Søndergaard, C. R.; Jensen, J. H. Graphical Analysis of PH-Dependent Properties of Proteins Predicted Using PROPKA. BMC Struct. Biol. 2011, 11, 6. https://doi.org/10.1186/1472-6807-11-6.

(42) Machado, M.; Pantano, S. Structural Bioinformatics SIRAH Tools: Mapping, Backmapping and Vis-Ualization of Coarse-Grained Models. Bioinformatics 2016, 32 (10), 2–3. https://doi.org/10.1093/bioinformatics/btw020.

(43) Abraham, M. J.; Murtola, T.; Schulz, R.; Páll, S.; Smith, J. C.; Hess, B.; Lindah, E. Gromacs: High Performance Molecular Simulations through Multi-Level Parallelism from Laptops to Supercomputers. SoftwareX 2015, 1-2, 19–25. https://doi.org/10.1016/j.softx.2015.06.001.

(44) Darre, L.; Machado, M. R.; Dans, P. D.; Herrera, F. E.; Pantano, S. Another Coarse Grain Model for Aquous Solvation: WAT FOUR? J. Chem. Theory Comput. 2010, 6 (12), 3793–07.

(45) Bussi, G.; Donadio, D.; Parrinello, M. Canonical Sampling through Velocity Rescaling. J. Chem. Phys. 2007, 126 (1), 014101. https://doi.org/10.1063/1.2408420.

(46) Parrinello, M.; Rahman, A. Polymorphic Transitions in Single Crystals: A New Molecular Dynamics Method. J. Appl. Phys. 1981, 52 (12), 7182–7190. https://doi.org/10.1063/1.328693.

(47) Heinig, M.; Frishman, D. STRIDE: A Web Server for Secondary Structure Assignment from Known Atomic Coordinates of Proteins. Nucleic Acids Res. 2004, 32 (WEB SERVER ISS.). https://doi.org/10.1093/nar/gkh429.

(48) Kuroda, Y.; Suenaga, A.; Sato, Y.; Kosuda, S.; Taiji, M. All-Atom Molecular Dynamics Analysis of Multi-Peptide Systems Reproduces Peptide Solubility in Line with Experimental Observations. Sci. Rep. 2016, 6 (December 2015), 1–6. https://doi.org/10.1038/srep19479.

